# Structural controllability predicts functional patterns and brain stimulation benefits associated with working memory

**DOI:** 10.1101/794388

**Authors:** L. Beynel, L. Deng, C.A. Crowell, M. Dannhauer, H. Palmer, S. Hilbig, A.V. Peterchev, B. Luber, S.H. Lisanby, R. Cabeza, L.G. Appelbaum, S.W. Davis

**Author notes:** Corresponding Author: Simon W Davis, PhD, 400 Trent Dr., Durham NC, 27710. These authors contributed equally to this work (L. Beynel, L. Deng).

## Abstract

The brain is an inherently dynamic system, and much work has focused on the ability to modify neural activity through both local perturbations and changes in the function of global network ensembles. Network controllability is a recent concept in network science that purports to predict the influence of individual cortical sites on global network states and state changes, thereby creating a unifying account of local influences on global brain dynamics. Here, we present an integrated set of multimodal brain–behavior relationships, acquired from functional magnetic resonance imaging during a transcranial magnetic stimulation intervention, that demonstrate how network controllability influences network function, as well as behavior. This work helps to outline a clear technique for integrating structural network topology and functional activity to predict the influence of a potential stimulation target on subsequent behaviors and prescribes next steps towards predicting neuromodulatory and behavioral responses after brain stimulation.

**Highlights:**

- This study tested the strength of network controllability using fMRI and rTMS
- Controllability correlates with functional modulation of working memory demand load
- Controllability is also correlated with the memory improvement from applied rTMS
- These findings link network control theory with physiology and behavior.

**In brief:** Beynel et al. show that the benefits of functionally targeted brain stimulation on working memory performance can be predicted by network control properties at the stimulated site. Structural controllability and functional activity independently predict this cognitive benefit.

**Author Contributions:** Conceptualization & Methodology: L.B, S.W.D., B.L., R.C., L.G.A.; Investigation: L.B., L.D., S.W.D., C.A.C., M.D., H.P., S.H.; Writing—Original Draft: L.B., L.D., S.W.D.; Writing—Review & Editing: L.B., L.D., S.W.D., L.G.A., A.V.P.; Funding Acquisition: S.W.D., R.C., B.L., S.H.L., A.V.P.; Resources: L.G.A., B.L., R.C.; Supervision: L.G.A., S.W.D.

## Introduction

A dominant narrative in contemporary neuroscience is that the brain is a vastly dynamic system of interacting networks called into action to achieve, maintain or change different mental states and accomplish cognition. Whereas previous views of neuroscience were largely reductionist or envisioned static topologies of connected communities, neuroscientists now consider cognition to result from controlled transitions between dynamically changing networks of neuronal ensembles. A key notion present in this network neuroscience view is that global brain states can be *controllable* from a minority of nodes, or even a single node (Betzel et al., 2016), thereby imparting certain privileged roles for different brain areas at certain times. Within this context, brain states can be altered through endogenous perturbations in these nodes that might result from changes in neuronal activity attributed to top-down attention, memory recollection or other cognitive processes (Kenett et al., 2018). Conversely, brain states can also be changed exogenously through the use of noninvasive neuromodulation approaches such as repetitive transcranial magnetic stimulation (rTMS). This latter approach is especially attractive because it raises the possibility to causally influence the brain in a controlled manner to test specific hypotheses about network dynamics.

While applications of control theory in the brain have gained substantial interest in the past few years, the validity of this framework has generated a lively theoretical debate (Pasqualetti et al., 2019; Tu et al., 2018), and questions remain about both the foundational tenets of the approach (e.g., *Are brain regions controllable?*) and the specific manner in which control is executed in the brain. Unfortunately, empirical evidence supporting the role of controllability in real-world applications is extremely limited. Recent theoretical and empirical work addressing neuronal control theory has focused on how the brain transitions from one state of activation to another (Betzel et al., 2016), and what anatomical substrates facilitate these transitions (Stiso et al., 2019). Within such a framework, *controllability* is the mathematical formulation of how a system can move through state-space along a desired trajectory (Gu et al., 2015). At its heart, network controllability theory assumes that the state of a system at a given time is a function of the previous state, the structural network linking the nodes (e.g. as measured through diffusion weighted imaging or DWI), and the additional energy perturbing it (Ruths and Ruths, 2014). In particular, of potential interest to the field of cognitive neuroscience is the concept of modal controllability (MC), a measure of the ability of a brain region to drive the brain as a network system into difficult-to-reach states (Medaglia et al., 2018; Pasqualetti et al., 2014). In light of these constructs, any satisfying account of MC would necessarily include the behavior generated by the system, and more importantly, (i) the functional activity that indexes brain states, and (ii) the controlled exogenous perturbation to manipulate them, both of which were the aims of investigation in the current study.

First, in order to understand how specific nodes can control brain state transitions, we utilize one of the most reliable brain patterns in fMRI-based cognitive neuroscience research: the increased activity in frontoparietal areas associated with greater working memory (WM) difficulty (Blumenfeld and Ranganath, 2007; Donner et al., 2002; Yeo et al., 2014). Within this network, prefrontal regions are generally associated with manipulation of information and response selection (D’Esposito et al., 1999; Postle, 2006), two cognitive processes associated with increasing difficulty. As noted above, MC has been inferred to quantify the capacity of a single node to drive the system to difficult-to-reach states. While this inference is driven largely by the frontoparietal distribution of MC (Gu et al., 2015), there is currently little concrete data supporting this interpretation (Tu et al., 2018). Nonetheless, several studies have examined the potential for MC to predict functional patterns through computer simulations (Betzel et al., 2016; Muldoon et al., 2016), but the identification of a reliable, significant association between MC and functional activity is still lacking (Cornblath et al., 2019), and a systematic investigation into the interdependencies between structural network controllability, functional activity, and cognition remains critical to address the value of this novel network control measure.

Second, in order to evaluate the value of MC in predicting dynamic state transitions, we used rTMS to test how the addition of exogenous stimulation affects brain states and behavior. While endogenous stimulation may be represented by a number of carefully controlled cognitive conditions (Cornblath et al., 2019; Medaglia et al., 2018), isolating the influence of one brain region in such approaches is difficult, if not impossible. rTMS therefore provides an ideal means to test the assumption that a localized change in energy to a modal controller could affect a global change in brain states and concomitant behavior. The effects of neuromodulation are typically observed not only in the stimulated site, but also in distal connected regions (Bestmann et al., 2004; Davis et al., 2017; Ruff et al., 2008; Wang et al., 2014; Wang et al., 2018), and thus the controllability framework offers an opportunity to probe the association between network properties and the effect of stimulation (Muldoon et al., 2016; Spiegler et al., 2016).

In the current analyses, we build on findings previously reported in Beynel et al (2019) and Davis et al (2018), to test the validity of network controllability by (i) testing whether MC tracked the empirically observed brain state transition as quantified by fMRI and (ii) if the MC of a site stimulated with rTMS predicted differences in behavioral changes during a concurrent WM task. To relate the theoretical to the actual state-transition in the brain, we used BOLD-related activity associated with increasing WM load. A higher parametric increase in BOLD-related activity suggests that additional neural resources can be relatively easy to recruit in more difficult task conditions, and thus indicates a relatively easy-to-reach brain state. Within this context, if MC represents the potential for a brain region to move from an easy to a difficult-to reach state, then we expect (i) *a negative linear relationship between MC and BOLD activity associated with increasing difficulty*. Next, if harder-to-reach states can be more easily achieved by adding exogenous stimulation with rTMS to a region with higher MC, then we expect (ii) *positive linear relationship between MC and rTMS effect*. Namely, subjects with higher modal controllability values in the stimulated site will show the greatest benefit from rTMS. If these hypotheses are confirmed, such findings will help validate the value of network controllability in predicting global brain states, and furthermore help predict rTMS effects that are foundational to a number of FDA-approved treatments.

## Results

In this 6-visit study, participants were screened, consented, and practiced a WM manipulation task (see Supplemental Methods) on Visit 1. On Visit 2 anatomical and diffusion-weighted imaging (DWI) scans were obtained as well as fMRI acquired while subjects performed the WM task with four difficulty levels (Very Easy, Easy, Medium, Hard) that were individualized based on performance from the first visit. These data were used to isolate a specific cortical target based on the BOLD response to the WM task (see overlap of stimulation sites in **Figure 1A**). On Visits 3 through 6 online, active or sham 5 Hz rTMS was applied while subjects performed the WM task at three difficulty levels (Easy, Medium, Hard).

**Figure 1.**
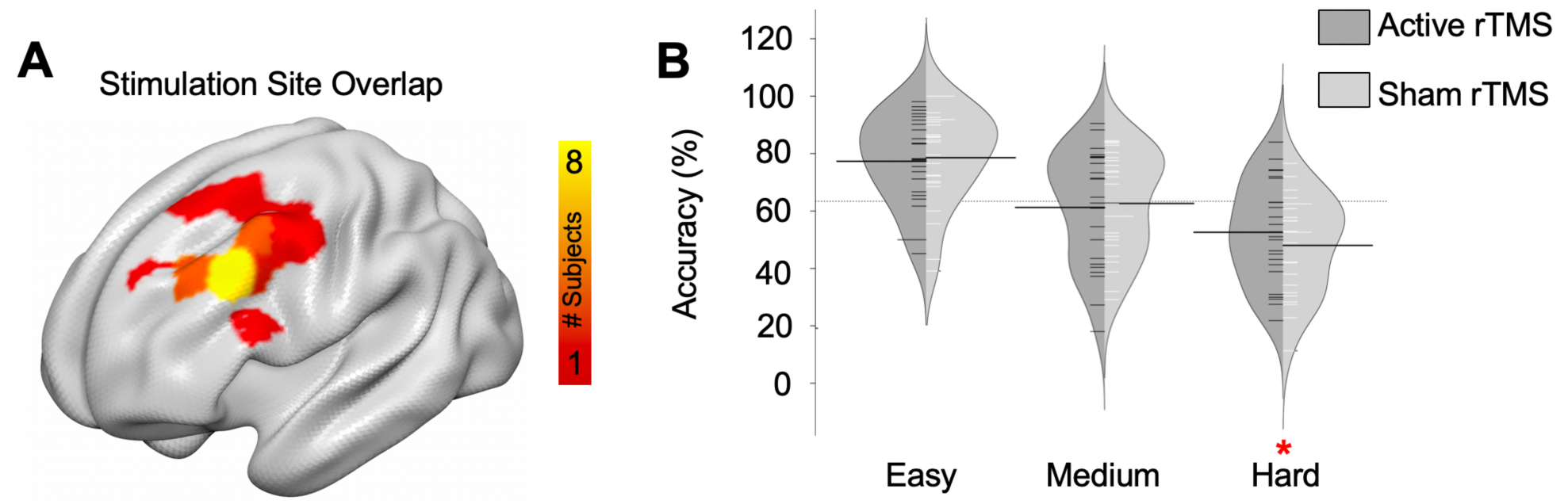
**A**: Overlap of stimulation targets for all participants, based on BOLD activity during a WM localizer task. B: Effects of online rTMS on WM accuracy at the group level.

A repeated measures ANOVA conducted on WM accuracy to evaluate effects of Difficulty (Easy, Medium, Hard) and Stimulation Type (Active, Sham) revealed a significant interaction between these two factors (F(2,44) = 3.04, p = 0.05). Decomposition of this interaction demonstrated that subjects were significantly more accurate with active, compare to sham rTMS, but only in the most difficult task condition (**Figure 1B**). Subsequent analyses used a within-subject metric to describe the percent increase in WM accuracy for active relative to sham rTMS, hereafter referred to as the “rTMS effect” (**Eq. 1**). More details on initial findings from this dataset can be found in Beynel et al. (2019) and Davis et al. (2018).

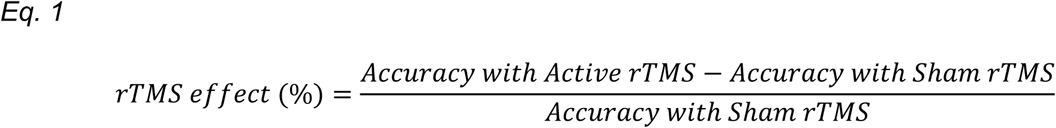

DWI-based streamline counts were used to construct structural networks for each participant, and MC values were computed across 471 cortical sites. **Figure 2A** illustrates the rank of group mean MC values for all regions across the brain (lighter warm colors reflecting greater MC). While the frontal regions stimulated in the current study (see inset and **Fig. 1**) were not selected on the basis of their MC value, the global distribution of MC values is nonetheless consistent with earlier investigations based on other cortical parcellations (Gu et al., 2015; Khambhati et al., 2018; Medaglia et al., 2018). Also consistent with Gu et al., the regions with higher MC were characterized by a lower structural connectivity strength with the rest of the brain (r_470_ = - 0.88; **Fig. 2C**). Univariate fMRI analysis revealed a typical pattern of BOLD activity associated with increasing WM load (lighter cool colors reflecting greater Functional Modulation, **Fig. 2B**). The spatial pattern of Functional Modulation was independent of the spatial pattern of MC, as evidenced by a weak correlation between the ranks of mean MC and mean Functional Modulation across regions (r_470_ = 0.11; **Fig. 2D**). Such independence between structural controllability and functional activity further suggests the possibility that the interplay between MC, Functional Modulation, and the behavioral effect of rTMS should be highly specific to the rTMS-targeted region.

**Figure 2.**
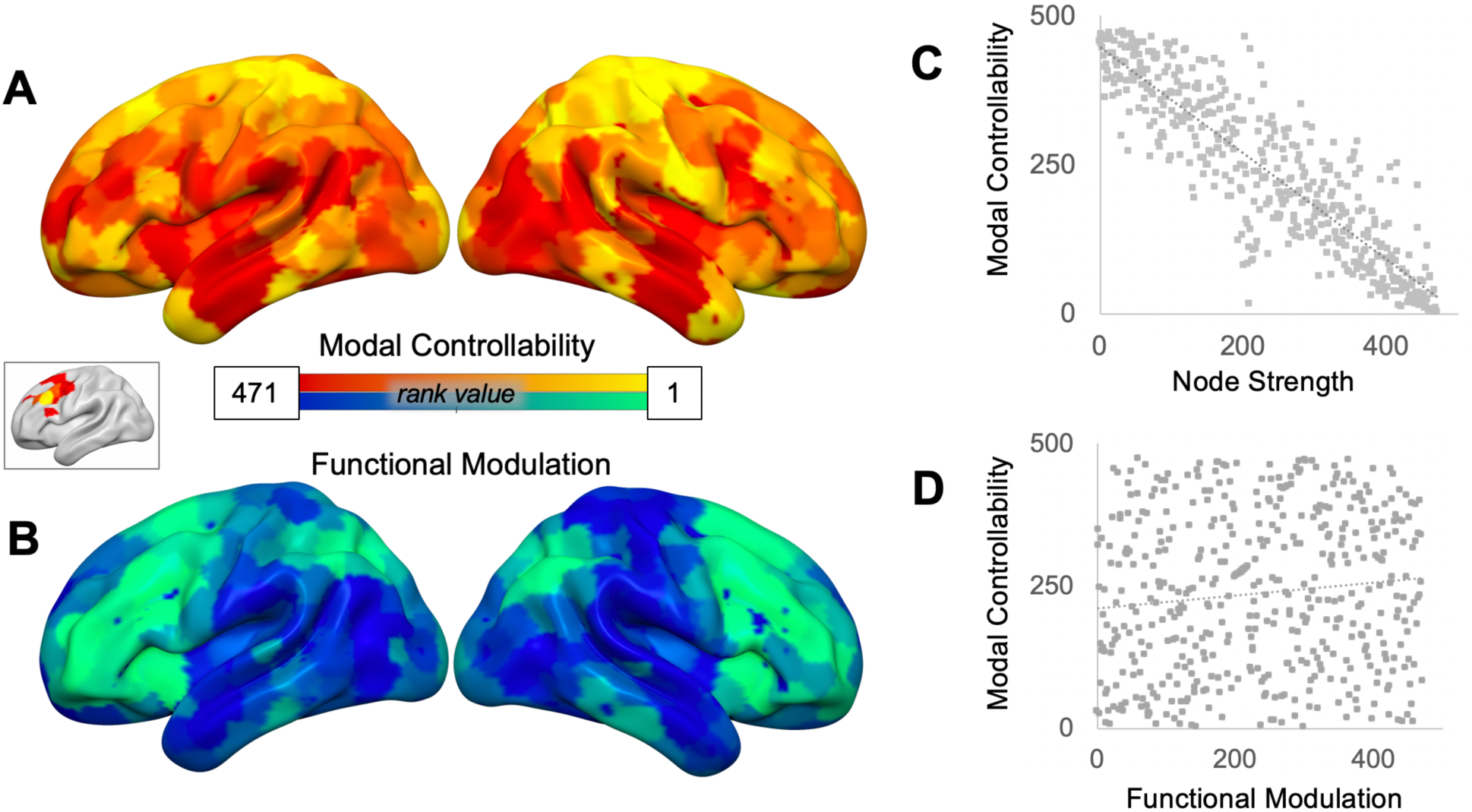
Mean modal controllability (**A**) and Functional Modulation (**B**) across all ROIs. Inset displays stimulation site overlap (see **Fig. 1**). As expected, MC showed a consistent linear relationship with node strength (**C**). While frontoparietal regions appear to show some overlap in both MC and Functional Modulation associated with increasing WM load, we observed no global relationship between these measures (**D**).

To test this hypothesis, we investigated these correlations at the targeted stimulation location and found that MC and fMRI-based Functional Modulation at the stimulated site were negatively correlated (**Fig. 3A**; r_22_ = - 0.53, p = 0.009), suggesting the co-occurrence of high MC and hard-to-reach brain state (i.e., weak Functional Modulation). Moreover, the average fMRI activation across these levels of difficulty was unrelated to MC (r_22_ = 0.04, p = 0.85), confirming that this result is indeed specific to task difficulty. Second, MC was significantly positively related (r_22_ = 0.46, p = 0.027) to the benefit from rTMS in difficult WM condition (**Fig. 3B**). This relationship was not present at Easy (r_22_ = 0.14, p = 0.51) or Medium (r_22_ = −0.054, p = 0.81) difficulty levels, confirming a selective rTMS effect at a difficult-to-reach cognitive state. Finally, the modulation of fMRI activation associated with difficulty at the stimulated ROI was also significantly related to the rTMS effect (**Fig. 3C**; r_22_ = −0.49, p = 0.017). This result supports the theory that that the effects of rTMS depend on brain state at the time of stimulation (Silvanto and Pascual-Leone, 2008), and suggests an interpretation that the benefit of rTMS is stronger when the stimulated region faces difficulty in effectively modulating its own state.

**Figure 3.**
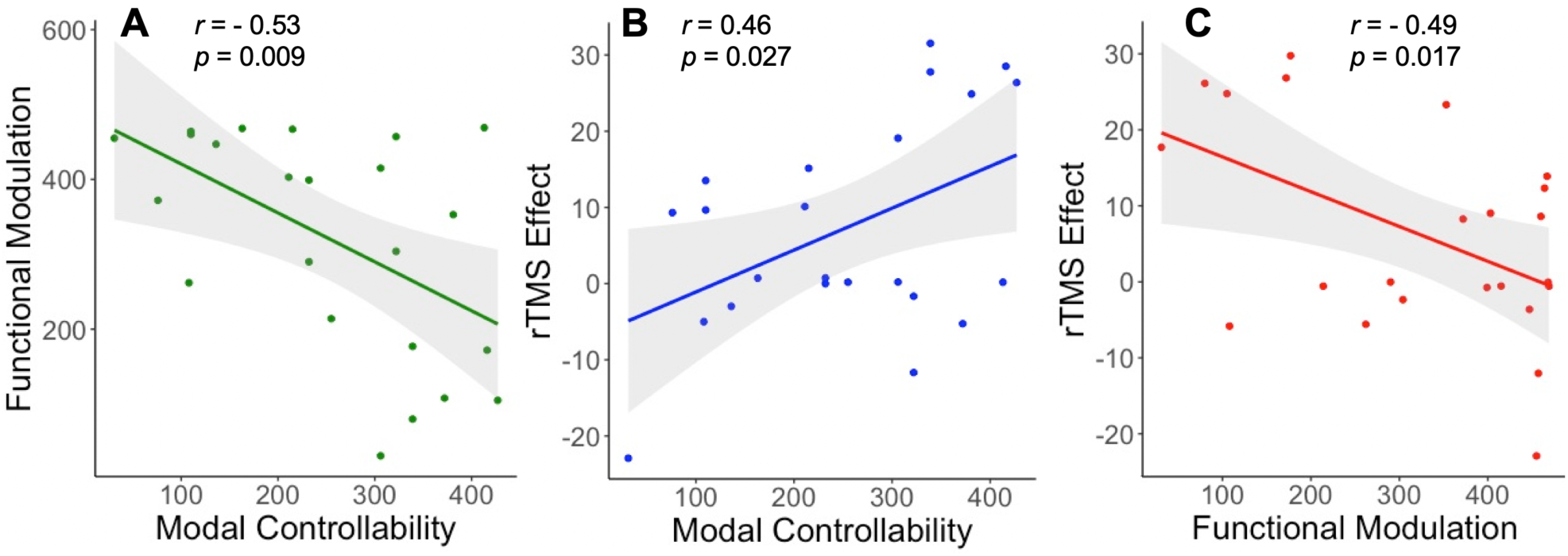
Relationship between Modal controllability and: **A**. Functional modulation, i.e. the parametric increase in BOLD-related activity associated with increasing working memory load; **B**. the rTMS effect, i.e. the percentage increase in WM accuracy after 5-Hz online rTMS. **C**: Correlation between Functional modulation and the rTMS effect. All correlations represent values derived from the stimulated ROI (See **Fig. 1**).

Finally, a mediation analysis was performed to test the hypothesis that the MC-rTMS relationship was mediated by the Functional Modulation associated with more difficult trials **(Fig. 4)**. This analysis yielded a significant total effect (TMS ∼ MC: z_c_ = 2.43, p = 0.03), as well as significant relationships between the predictor and mediator (TMS ∼ Functional Modulation: z_a_ = - 3.36, p = 0.0008) and the mediator and outcome variable (Functional Modulation ∼ MC: z_b_ = - 2.69, p < 0.007). Crucially, a significant mediation by Functional Modulation (z_ab_ = 2.12, p = 0.04) explained a major proportion of the total effect (64%), resulting in a non-significant direct contribution of the rTMS effect to MC (z_c’_ = 0.68, p = 0.53). To ensure the reliability and specificity of the mediation by Functional Modulation, we repeated the analysis on all possible alternative mediation models, where the directions of mediation and the mediator were changed. Here, no strong mediation effects were observed in these alternative models (see Supplementary Information), further confirming that the observations in **Figure 2** are best explained as functional activity mediating the relationship between rTMS effect and structural controllability.

**Figure 4.**
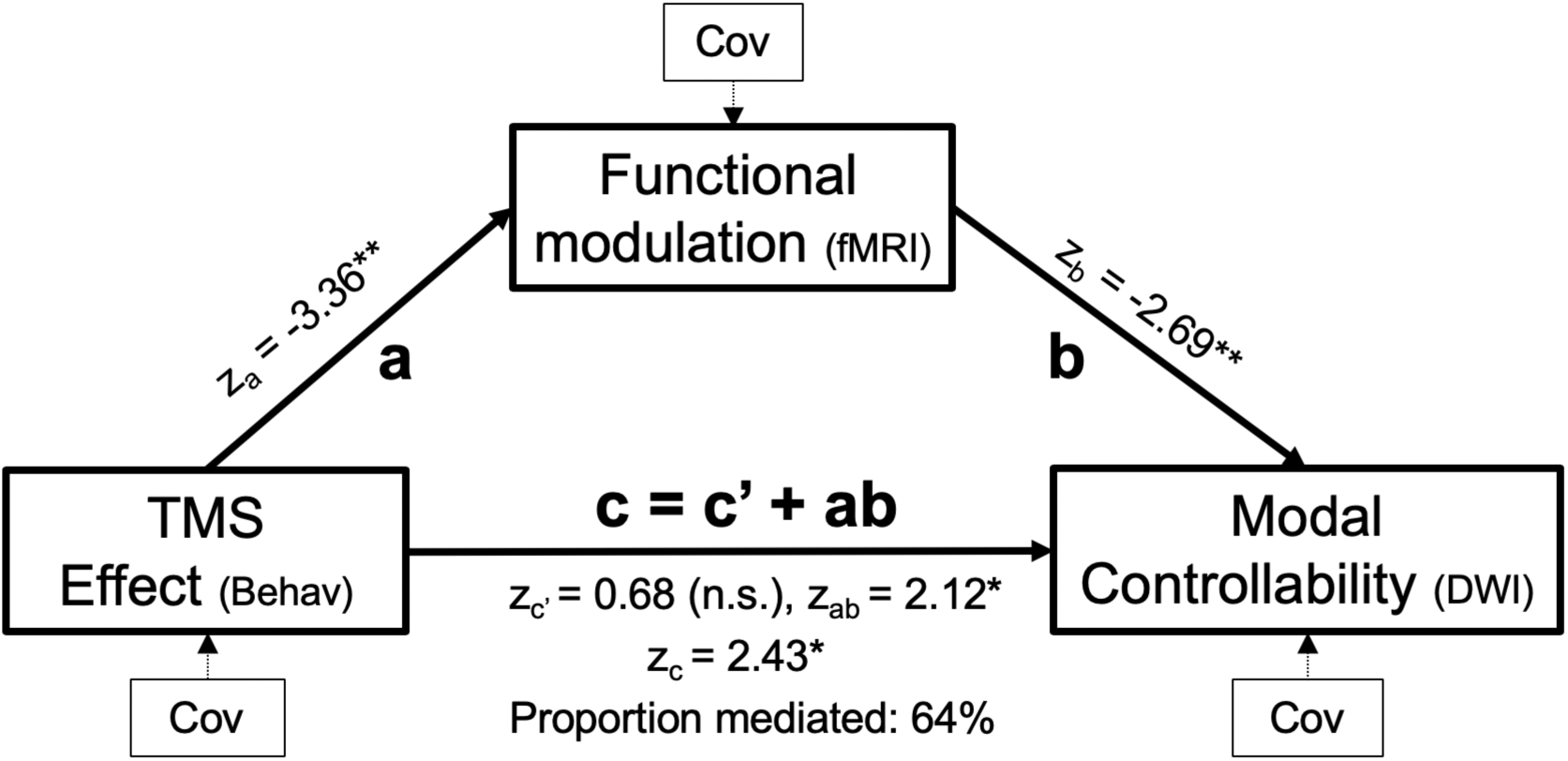
Mediation analysis investigating the capacity for fMRI activation to explain the relationship between Modal Controllability and rTMS effect. Age, gender, and baseline task performance were controlled as confounding covariates (Cov).

## Discussion

The current study sought to inform the validity of network controllability in characterizing the ability of a single cortical node to guide the trajectory of global brain states. We used DWI and fMRI to investigate the functional and structural information associated with brain state transitions, and rTMS to test whether providing exogenous energy to a specific node could help the brain transition to difficult-to-reach states. The notion that brain activity could be controlled by energy input to a single cortical node is still highly debated (Pasqualetti et al., 2019; Tu et al., 2018), and only a handful *in vivo* studies have investigated its validity in human models (Medaglia et al., 2018; Stiso et al., 2019). Here we examined the interactions between controllability, brain states, neuromodulation and behavior and found significant relationships between these measures, which can be interpreted as the cost associated with transitions needed to reach a cognitive state as resolved in our single mediation model. These results help to show that brain network information, based on control theory, is supported by both functional and neuromodulatory data, and helps support the use of a new family of targeting approaches based on the dynamic control of global brain states.

A principle finding from this study was the observation that MC was reliably associated with the functional modulation of BOLD activity associated with parametric increases in WM load (**Fig. 3A**). Moreover, it was found that MC at the stimulated site was reliably predicted by the benefit from rTMS in the WM task. This rTMS benefit was only found in the most difficult condition (**Fig. 1B**), and the association between MC and this performance benefit was selective to this difficult condition (**Fig. 3B**). This result, therefore, supports existing computational models of network dynamics based on structural connectivity information, which purport to describe the energy associated with transitioning between brain states. Predicting the effect of brain stimulation by utilizing brain network topology is of theoretical and applied significance, and the current results provide a mechanistic application of these principles as set forth by Pasqualetti et al. (2014). According to these authors, the brain constitutes a controllable network that can be moved into different states with input perturbation, such as brain stimulation. A recent study provided behavioral evidence supporting theory that the controllability is a factor guiding the influence of brain stimulation (Medaglia et al., 2018); the authors observed that a median split participants based on MC values identified subtle performance differences in a language production task, providing some support for the conclusion that MC may influence task-based response to rTMS. Nonetheless, the observed effects did not demonstrate evidence that higher MC allows the brain to more easily traverse difficult-to-reach states, as the influence of modal controllability was found only in the easiest language task condition. Furthermore, this condition did not show a significant benefit of active rTMS over sham and so it is difficult to draw conclusions of the functional relevance of MC in that study.

Finally, we found that the core relationship between rTMS effect and MC was significantly mediated by the functional modulation associated with increasing difficulty. The mediation observed above extends this inference to suggest MC reflects not only the structural value, but also the *functional* capacity for a given node to modify the derived eigenmatrix. This result helps to advance the notion that all three metrics examined here—the behavioral benefit from rTMS, Functional Modulation, and MC—reflect distinct constructs of “difficulty”. In the current approach, we posited that the networks identified by parametric changes in functional activation can be influenced by modulation of a single site. Following Gu et al. (2015), we treated this influence as an eigenvalue problem, such that regions with high MC have the greatest influence on less persistent eigenmodes. Thus, MC herein reflects the ability to drive the dynamics of a network towards hard-to-reach configurations. This mathematical formulation helps to explain the more basic mediation finding, in that while MC of the network reflects the *capacity* for a node to shift to a new brain state, Functional Modulation reflects the degree to which regions effectively *do traverse* such a brain state. These cooperative contributions in turn leads to greater performance improvements after rTMS, and presumably a successful shift in brain state. Even so, we agree with Tu et al. (2018) that this is an interpretation at the level of *regions* and may not capture multivariate interactions inherent in the broader question, *Is the brain controllable?* The current analysis nonetheless seeks to ground this interpretation by applying the control theory metrics to real human brain data.

The approach used here has clear implications for the individualization of TMS targeting, and advances targeting approaches that have improved from scalp-based, to anatomical, to functionally-guided techniques (Herwig et al., 2001). Both clinical and basic science targeting approaches for TMS should consider the consequences of regional brain stimulation in manipulating more global functional and structural networks connected to the stimulation site. When regional stimulation is applied to models that differ only in their structural connectivity, the resulting activity patterns differ between individuals (Jbabdi et al., 2015), suggesting that functional responses are sensitive to subject-specific differences in white matter structure. Computational models informed by both forms of brain information are therefore necessary to provide realistic, individualized models of immediate TMS outcomes. Our findings relating the structural and functional components of a controllable system suggests that modal controllability may be used as an alternative when functional localizers for TMS are unfeasible.

In sum, by combining information from structural network topology, whole-brain functional activity associated with a WM task, and brain stimulation, we confirmed the validity of network controllability by showing a strong interdependence of these measures. This work helps to outline a clear technique for integrating structural network topology and functional activity to inform network-based approaches in selecting an individualized specific cortical target for neurostimulation that could help increasing rTMS efficacy.

## Material and Methods

The following sections give a brief overview of the methods, additional information can be found in Beynel et al. (2019), and in the Supplementary Materials.

### Participants

Twenty-nine young adults (mean 23.38 ± 5.13 years old, 14 females) completed this 6-day study, approved by the Duke Medical School IRB (#Pro00065334); 5 subjects were subsequently removed due to incomplete imaging data. We used a modified Sternberg task in which arrays of letters were manipulated within WM (D’Esposito et al., 1999), noted here as the delayed-response alphabetization task (DRAT). For each trial, letters were presented, followed by a delay period during which participants mentally reorganized letters into alphabetical order. A letter with a number above it was then presented, and participants were asked whether the letter was (1) New trials not in the original set, (2) Valid trials comprising a letter from the original array, and the number matched the serial position of the letter once the sequence was alphabetized, or (3) Invalid trials, where the probe was in the original array but the number did not match the serial position of the alphabetized probe.

After an initial screening visit, participants were scanned in a functional localizer MRI session at a 3-T gradient-echo scanner (General Electric 3.0 Tesla Sigma Excite HD short bore scanner), equipped with an 8-channel head coil. During this session, a structural MRI and a diffusion weighted imaging (DWI) scan were acquired, as well as 4 blocks of functional acquisitions while participants performed the DRAT. fMRI activations were defined using an event-related design with the array, the delay, and the response periods. A parametric regressor was used for the delay period, reflecting increases in brain activation associated with increases in task difficulty, as defined by load (i.e., the number of letters in an array). The peak activation within the left medial frontal gyrus in each participant was chosen as the rTMS target and entered into the neuronavigation system (BrainSight, Rogue Research, Canada). Sham stimulation was applied using the same coil in placebo mode, which produced clicking sounds and somatosensory sensation via electrical stimulation with scalp electrodes mimicking those occurring in the active mode, but without a significant E-field induced in the brain (Smith, & Peterchev, 2018).

### Controllability Measures

We adopted a mathematical definition of controllability from Pasqualetti et al. (2014) where the brain system dynamics is modelled with a linear discrete-time and time-invariant model:

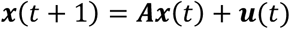

Here, the 471-by-471 weighted adjacency matrix ***A*** describes the structural connectivity between each pair of nodes. The 471-by-1 column vector ***x***(*t*) describes the brain state. The 471-by-1 column vector ***u***(*t*) indicates external input, where the only non-zero entry *u*_*i*_(*t*) denotes TMS stimulation at brain region *i*. In the current analysis we focus on modal controllability (Kalman, 1964), interpreted here and elsewhere (Gu et al., 2015) as the ability of one region to steer the brain towards hard-to-reach states. The modal controllability of brain region *i* is defined thus as

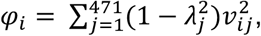

where *λ*_*j*_ denotes the *j-th* eigenvalue of the network adjacency matrix ***A***, and *v*_*ij*_ denotes the *i-th* entry in *λ*_*j*_’s corresponding eigenvector ***v***_*j*_. The eigenvectors ***v***_*i*_ represent the ‘mode states’ of the brain network system, whose linear combination can potentially describe any given activation pattern of the brain. The eigenvalues *λ*_*i*_ can be interpreted as the persistence of the corresponding mode states, thus a small *λ* is indicative of a transient, hard-to-reach state. By extension of the Popov-Belovich-Hautus test (Lathi, 2005), the controllability of the *j-th* mode in region *i* is proportional to *v*_*ij*_. Therefore, as a composite measurement, the modal controllability theoretically predicts whether stimulating a brain region can move the brain into hard-to-reach states *in general*. Though current methodology impedes a prediction about which specific hard-to-reach state the brain has transitioned to, here we rely on the brain state pattern associated with increasing difficulty WM trials. Rank was used instead of the actual value to control for cross-subject noise.

## Acknowledgments

The authors wish to thank Duy Nguyen, Alexandra Brito, and Joyce Wang for assistance with data collection.

## References

Bestmann, S., Baudewig, J., Siebner, H.R., Rothwell, J.C., and Frahm, J. (2004). Functional MRI of the immediate impact of transcranial magnetic stimulation on cortical and subcortical motor circuits. Eur J Neurosci 19, 1950–1962.

Betzel, R.F., Gu, S., Medaglia, J.D., Pasqualetti, F., and Bassett, D.S. (2016). Optimally controlling the human connectome: the role of network topology. Sci Rep 6, 30770.

Beynel, L., Davis, S., Crowell, C., Hilbig, S., Lim, W., Nguyen, D., Palmer, H., Brito, A., Peterchev, A., and Luber, B. (2019). Online repetitive transcranial magnetic stimulation during working memory in younger and older adults: A randomized within-subject comparison. PloS one 14, e0213707.

Blumenfeld, R.S., and Ranganath, C. (2007). Prefrontal cortex and long-term memory encoding: an integrative review of findings from neuropsychology and neuroimaging. The Neuroscientist 13, 280–291.

Cornblath, E.J., Tang, E., Baum, G.L., Moore, T.M., Adebimpe, A., Roalf, D.R., Gur, R.C., Gur, R.E., Pasqualetti, F., Satterthwaite, T.D., and Bassett, D.S. (2019). Sex differences in network controllability as a predictor of executive function in youth. NeuroImage 188, 122–134.

D’Esposito, M., Postle, B.R., Ballard, D., and Lease, J. (1999). Maintenance versus manipulation of information held in working memory: An Event-Related fMRI study. Brain and cognition 41, 66–86.

Davis, S.W., Luber, B., Murphy, D.L.K., Lisanby, S.H., and Cabeza, R. (2017). Frequency-specific neuromodulation of local and distant connectivity in aging and episodic memory function. Hum Brain Mapp.

Donner, T.H., Kettermann, A., Diesch, E., Ostendorf, F., Villringer, A., and Brandt, S.A. (2002). Visual feature and conjunction searches of equal difficulty engage only partially overlapping frontoparietal networks. NeuroImage 15, 16–25.

Gu, S., Pasqualetti, F., Cieslak, M., Telesford, Q.K., Yu, A.B., Kahn, A.E., Medaglia, J.D., Vettel, J.M., Miller, M.B., Grafton, S.T., and Bassett, D.S. (2015). Controllability of structural brain networks. Nat Commun 6, 8414.

Herwig, U., Schonfeldt-Lecuona, C., Wunderlich, A.P., von Tiesenhausen, C., Thielscher, A., Walter, H., and Spitzer, M. (2001). The navigation of transcranial magnetic stimulation. Psychiatry Res 108, 123–131.

Jbabdi, S., Sotiropoulos, S.N., Haber, S.N., Van Essen, D.C., and Behrens, T.E. (2015). Measuring macroscopic brain connections in vivo. Nat Neurosci 18, 1546–1555.

Kalman, R.E. (1964). When is a linear control system optimal? Trans ASME J of Basic Engineering, Ser D

Kenett, Y.N., Medaglia, J.D., Beaty, R.E., Chen, Q., Betzel, R.F., Thompson-Schill, S.L., and Qiu, J. (2018). Driving the brain towards creativity and intelligence: A network control theory analysis. Neuropsychologia 118, 79–90.

Khambhati, A.N., Medaglia, J.D., Karuza, E.A., Thompson-Schill, S.L., and Bassett, D.S. (2018). Subgraphs of functional brain networks identify dynamical constraints of cognitive control. PLoS computational biology 14, e1006234.

Lathi, B.P. (2005). Linear systems and signals, 2nd edn (New York: Oxford University Press).

Medaglia, J.D., Harvey, D.Y., White, N., Kelkar, A., Zimmerman, J., Bassett, D.S., and Hamilton, R.H. (2018). Network Controllability in the Inferior Frontal Gyrus Relates to Controlled Language Variability and Susceptibility to TMS. J Neurosci 38, 6399–6410.

Muldoon, S.F., Pasqualetti, F., Gu, S., Cieslak, M., Grafton, S.T., Vettel, J.M., and Bassett, D.S. (2016). Stimulation-Based Control of Dynamic Brain Networks. PLoS computational biology 12, e1005076.

Pasqualetti, F., Gu, S., and Bassett, D.S. (2019). RE: Warnings and caveats in brain controllability. NeuroImage 197, 586–588.

Pasqualetti, F., Zampieri, S., and Bullo, F. (2014). Controllability metrics, limitations and algorithms for complex networks. IEEE Trans Control Netw Syst 1, 40–52.

Postle, B.R. (2006). Working memory as an emergent property of the mind and brain. Neuroscience 139, 23–38.

Ruff, C.C., Bestmann, S., Blankenburg, F., Bjoertomt, O., Josephs, O., Weiskopf, N., Deichmann, R., and Driver, J. (2008). Distinct causal influences of parietal versus frontal areas on human visual cortex: evidence from concurrent TMS-fMRI. Cerebral cortex 18, 817–827.

Ruths, J., and Ruths, D. (2014). Control profiles of complex networks. Science 343, 1373-1376.

Silvanto, J., and Pascual-Leone, A. (2008). State-dependency of transcranial magnetic stimulation. Brain Topogr 21, 1–10.

Spiegler, A., Hansen, E.C., Bernard, C., McIntosh, A.R., and Jirsa, V.K. (2016). Selective Activation of Resting-State Networks following Focal Stimulation in a Connectome-Based Network Model of the Human Brain. eNeuro 3.

Stiso, J., Khambhati, A.N., Menara, T., Kahn, A.E., Stein, J.M., Das, S.R., Gorniak, R., Tracy, J., Litt, B., Davis, K.A., et al. (2019). White Matter Network Architecture Guides Direct Electrical Stimulation through Optimal State Transitions. Cell Rep 28, 2554–2566 e2557.

Tu, C., Rocha, R.P., Corbetta, M., Zampieri, S., Zorzi, M., and Suweis, S. (2018). Warnings and caveats in brain controllability. NeuroImage 176, 83–91.

Wang, J.X., Rogers, L.M., Gross, E.Z., Ryals, A.J., Dokucu, M.E., Brandstatt, K.L., Hermiller, M.S., and Voss, J.L. (2014). Targeted enhancement of cortical-hippocampal brain networks and associative memory. Science 345, 1054–1057.

Wang, W.C., Wing, E.A., Murphy, D.L.K., Luber, B.M., Lisanby, S.H., Cabeza, R., and Davis, S.W. (2018). Excitatory TMS modulates memory representations. Cogn Neurosci 9, 151–166.

Yeo, B.T., Krienen, F.M., Eickhoff, S.B., Yaakub, S.N., Fox, P.T., Buckner, R.L., Asplund, C.L., and Chee, M.W. (2014). Functional Specialization and Flexibility in Human Association Cortex. Cerebral cortex.

