## Supplemental Text and Figures for "Structural controllability predicts functional patterns and brain stimulation benefits associated with working memory"

Supplementary Material

The following sections first provide a brief overview of the study, followed by additional information regarding data processing that allows calculating network controllability.

### Study overview: Experimental design

Twenty-nine healthy young adults performed a 6-day protocol (**Figure S1**); 5 subjects were subsequently removed due to incomplete imaging data. During the first visit, subjects consented to participate, were screened in to make sure they did not have any contraindications to TMS or MRI. Resting motor threshold was then assessed and participants performed the working memory (WM) task. During the second visit, MRI was performed in a 3-T GE scanner at the Duke Brain Imaging Analysis Center (BIAC). Structural MRI and diffusion weighted imaging (DWI) were followed by 4 fMRI runs during which participants performed the WM task. The anatomical MRI was acquired using a 3D T1-weighted echo-planar sequence (matrix = 2562, time repetition [TR] = 12 ms, time echo [TE] = 5 ms, field of view [FOV] = 24 cm, slices = 68, slice thickness = 1.9 mm, sections = 248). DWI data were collected using a single-shot echo-planar imaging sequence (TR = 1700 ms, slices = 50, thickness = 2.0 mm, FOV = 256 × 256 mm2, matrix size 128 × 128, voxel size = 2 mm3, b-value = 1000 s/mm2, diffusion-sensitizing directions = 36, total images = 960, total scan time = 5 min). Finally, in the fMRI runs, coplanar functional images were acquired using an inverse spiral sequence (64 × 64 matrix, TR = 2000 ms, TE = 31 ms, FOV = 240 mm,

37 slices, 3.8-mm slice thickness, 254 images). The total scan time was approximately 1 h 40 min. fMRI analysis was then performed (see fMRI analysis section) in order to define a target for the subsequent rTMS visits. During visits 3-6, rTMS was performed with an active/placebo figure-8 coil (A/P Cool-B65) and a MagPro X100 stimulator with MagOption (MagVenture, Denmark). The coil position was continually monitored through a stereotaxic neuronavigation system (Brainsight, Rogue Research, Canada) and maintained at a high level of precision throughout the session with real-time robotic guidance (Smart Move Robot, Advanced Neuro Technology, Netherlands).

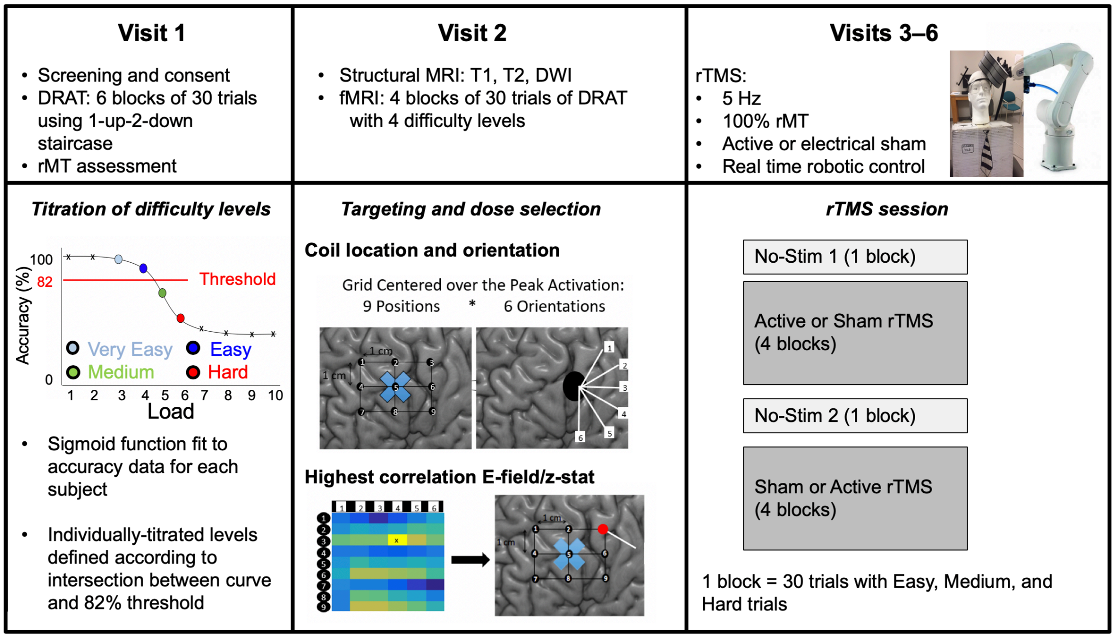


*__Figure S1__: Schematic representation of the full protocol.*

### __fMRI analysis__

Functional images were preprocessed using image processing tools, including FLIRT (FMRIB's Linear Image Registration Tool) and FEAT (FMRIB Expert Analysis Tool) from FMRIB's Software Library (FSL, http://fmrib.ox.ac.uk/fsl). Images were corrected for slice acquisition timing, motion, and linear trend; motion correction was performed using FSL's MCFLIRT, and 6 motion parameters estimated from the step were then regressed out of each functional voxel using standard linear regression. Images were then temporally smoothed with a high-pass filter using a 190 s cutoff, and normalized to the Montreal Neurological Institute (MNI) stereotaxic space. White matter and CSF signals were also removed from the data and regressed from the functional data using the same method as the motion parameters. Spatial filtering with a Gaussian kernel of full-width half-maximum (FWHM) of 6 mm was applied. BOLD activations during the array presentation, the delay period, and the response period were entered in a standard general linear model (GLM), using HRF-convolved trial regressors and the temporal derivatives. Importantly, the Functional Modulation (FM) of task difficulty was modelled by an additional parametric regressor and its temporal derivative during the delay period, reflecting the parametric increase in brain activation associated with the increase in task difficulty. These parametric regressors were orthogonalized with the delay period regressor. Since the delay period has been associated with working memory process, the delay period parametric activation was chosen as a measure of working memory state transition. For TMS targeting, the peak activation within left MFG was selected in each participant, and entered into the neuronavigation system (BrainSight, Rogue Research, Canada).

To determine the TMS-induced electric field (E-field), simulations using T1, T2, and DWI images and the finite element method in the SimNIBS software package [ref] were conducted. The models featured five distinct tissue types: skin, skull, cerebrospinal fluid (CSF), gray matter, and white matter. The DWI information was used to generate anisotropic conductivities for white matter using the volume-normalized approach. E-field was simulated at 54 candidate coil placements: 9 positions generated by placing a 3 × 3 grid with 1 $cm^2$ spacing above the peak fMRI activation; and 6 orientations per position corresponding to 30° rotation increments in a 180° semicircle. The coil position and orientation showing the highest correlation with fMRI activation pattern was selected as the rTMS target.

### Data processing for network controllability

#### Structural connectivity

Information on the structural connections based on diffusion tractography, between each pair of regions in our data were assessed with a standard DWI processing pipeline used previously in our group. DWI data were analyzed utilizing FSL (https://fsl.fmrib.ox.ac.uk/fsl/fslwiki) and MRtrix (http://mrtrix.org) software packages. Data were de-noised, corrected with eddy current correction, and bias-field corrected using MRtrix (Tournier et al., 2007) and FSL. Constrained spherical deconvolution (CSD) was utilized in calculating the fiber orientation distribution (FOD). This FOD was used along with the brain mask to generate whole brain tractography. Relevant parameters regarding track generation were as follows: seed = at random within mask; step-size = 0.2 mm; 10,000,000 tracts. After tracts were generated, they were filtered using spherical-deconvolution informed filtering of tractograms (Smith et al., 2012) in order to improve the quantitative nature of the whole-brain streamline reconstructions used here. This process utilizes an algorithm which determines whether a streamline should be removed or not based off of information obtained from the FOD, which improves the selectivity of structural connectomes by using a cost-function to eliminate false positive tracts. Tracts were SIFTed until 1 million tracts remained. Connectomes were then generated by using FLIRT to apply a linear registration to the HOA atlases mentioned above to register them to native diffusion space; subsequent connectomes describe the number of streamlines connecting any pair of regions within the HOA atlas.

#### Cortical Parcellation and ROI extraction

A consistent parcellation scheme across all subjects and all modalities (DWI, fMRI) was used. Subjects' T1-weighted images were segmented using SPM12 (www.fil.ion.ucl.ac.uk/spm/software/spm12/), yielding a grey matter (GM) and white matter mask in the T1 native space for each subject. The entire GM was then parcellated into 471 regions of interest (ROIs), each representing a network node by using a sub-parcellated version of the Harvard-Oxford Atlas, defined originally in MNI space. The T1-weighted image was then nonlinearly normalized to the ICBM152 template in MNI space using fMRIB's Non-linear Image Registration Tool (FNIRT, FSL, www.fmrib.ox.ac.uk/fsl/). The inverse transformations were applied to the HOA atlas in the MNI space, resulting in native-T1-space GM parcellations for each subject. Then, T1-weighted images were co-registered to native diffusion space using the subjects' unweighted diffusion image as a target; this transformation matrix was then applied to the GM parcellations above, using FSL's FLIRT linear registration tool, resulting in a native-diffusion-space parcellation for each subject. For each subject, the peak fMRI activations coordinates were transformed back into MNI coordinates. MRICron was then used by adding the 471 ROIs on an MNI brain, and the ROI number corresponding to the peak coordinates was then extracted as the "individualized stimulated site ROI", the same ROI number was used to define the modal controllability values at the targeted site.

### Statistical analysis

Mediation analysis was performed with structural equation modelling (SEM), using the *lavaan* package (version 0.5, Rosseel, 2012) in R version 3.3.3 (R Development Core Team, 2016). The standard errors in the mediation model were estimated using robust SEM option. In addition, age, gender, and starting set size are modeled as three independent covariates to eliminate potential confounding effect to each of the three variables of interest, i.e., TMS effect, modal controllability, and Functional Modulation.

To ensure the reliability of the mediation analysis, we repeated the analysis on all possible alternative mediation models, where the directions of mediation and the mediator were reversed. As is shown in **Figure S2**, among all alternative mediation models, only the first model with Functional Modulation as the mediator between MC and TMS showed a marginally significant mediation effect accompanied by a considerably large proportion (63%) of total effect being mediated.

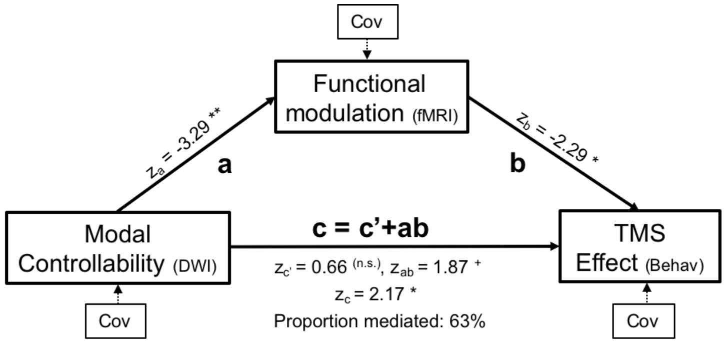


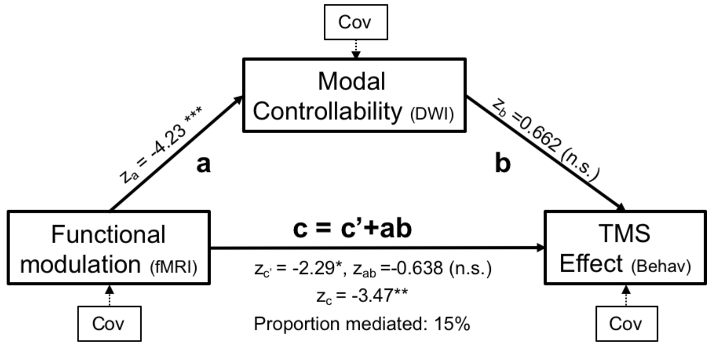


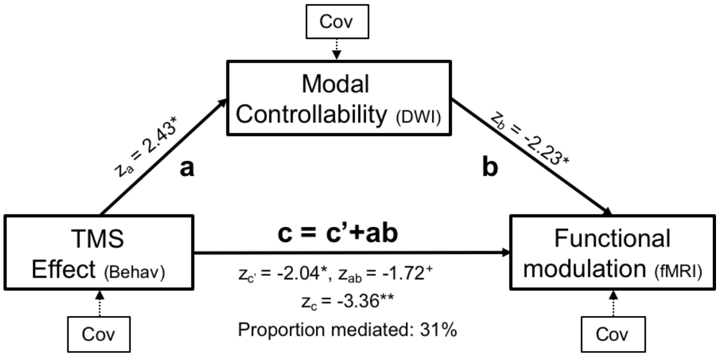


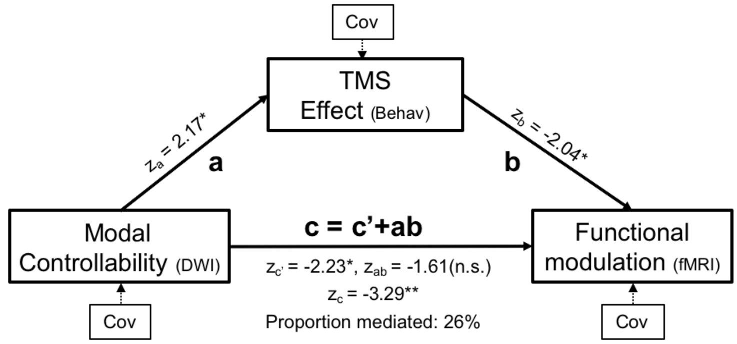

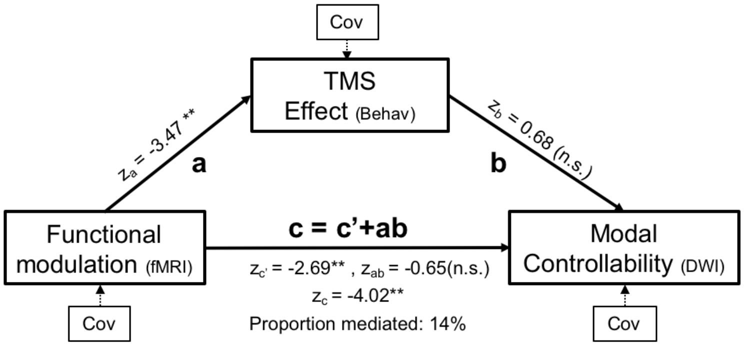


*Figure S2: Alternative mediation models and their corresponding statistical values.*

### References

Rosseel, Y. (2012). lavaan: An R Package for Structural Equation Modeling. J Stat Softw 48, 1-36.

Smith, R.E., Tournier, J.D., Calamante, F., and Connelly, A. (2012). Anatomically-constrained tractography: improved diffusion MRI streamlines tractography through effective use of anatomical information. Neuroimage 62, 1924-1938.

Tournier, J.D., Calamante, F., and Connelly, A. (2007). Robust determination of the fibre orientation distribution in diffusion MRI: non-negativity constrained super-resolved spherical deconvolution. Neuroimage 35, 1459-1472.
